# Untangling mechanisms for cerebellar neural specification using human pluripotent stem cell-derived organoids

**DOI:** 10.64898/2026.04.27.720597

**Authors:** Sergio Helgueta, Antonela Bonafina, Nicolas Olivié, Bernard Coumans, Laurent Nguyen, Ira Espuny-Camacho

## Abstract

The cerebellum is one of the most complex structures of the brain composed of a high diversity of GABAergic and glutamatergic neurons. Whereas cerebellar biogenesis has been extensively studied in the mouse, an in-depth characterization of genes and pathways involved in cerebellar specification and maturation in the humans remains overlooked. Here, we used human pluripotent stem cells (hPSC)-derived cerebellar organoids (CRBOs) to study the temporal biogenesis of neuronal subtypes. Our results show that CRBOs acquire caudal neural tube identity at an early stage followed by a time-dependent expression of mature cerebellar neuronal markers *in vitro*, mimicking human neurodevelopment. CRBOs show the generation of both cerebellar excitatory and inhibitory neurons and the expression of glial cell markers, suggesting the generation of a high variety of cerebellar cell types *in vitro*. Further, *in vitro* CRBOs show expression of cerebellar disease associated genes, such as those related to ataxia. Our results establish CRBOs as a valuable platform to explore the mechanisms of human cerebellar development and related disorders.

## Introduction

The cerebellum is the largest structure of the hindbrain, located in the posterior cranial fossa (D. C. Van Essen et al., 2018) and associated to diverse cerebellar-specific functions (Stoodley et al., 2012). The development of the cerebellum is governed by the tight coordinated action of transcription factors, such as GBX2, EN2 and PAX2 and extrinsic factors and signaling pathways, such as fibroblast growth factors (FGFs) and sonic hedgehog (SHH) signaling, that regulate its specification and patterning (Butts et al., 2014; M. J. van Essen et al., 2020). While classically associated with motor coordination, balance, posture, and motor learning, the cerebellum also contributes to higher-order functions, including cognition, social behavior, and language processing (Cutando et al., 2022; Marek et al., 2018; Schmahmann, 2019). Cerebellar biogenesis entails the production of diverse inhibitory GABAergic subtypes – including Purkinje cells (PCs), nucleo-olivary projection neurons, and various interneurons – alongside excitatory glutamatergic populations such as granule cells (GCs), unipolar brush cells and deep cerebellar nuclei (DCN) neurons (Akazawa et al., 1995; Ben-Arie et al., 1997; Hoshino et al., 2005; MacHold & Fishell, 2005; Pascual et al., 2007; V. Y. Wang et al., 2005). These lineages originate from two progenitor zones in the cerebellum, the PTF1A+, KIRREL2+ ventricular zone (VZ) region and the ATOH1+ rhombic lip (RL), which generate GABAergic and glutamatergic progenies, respectively (Marzban et al., 2015). At birth, RL progenitors migrate to establish the external granular layer (EGL) where they continue to divide before exiting cell cycle to produce GCs. Newly generated GCs migrate from the EGL, along Bergmann glia and across the PC layer, to settle in the internal granular layer (IGL). Key signals regulating this migration include SHH, which is secreted by PCs to promote proliferation of GC precursors, and Reelin, secreted by Cajal-Retzius cells which controls migration and organization of both PCs and GCs within the IG (D’Arcangelo, 2014; Miyata et al., 2010).

Glutamatergic GCs are the most abundant cell type within the cerebellum, and further within the brain, whereas PCs constitute the sole output neurons of the cerebellar cortex (Fatemi et al., 2013; Mapelli et al., 2022; Sathyanesan et al., 2019). Cerebellar pathologies encompass a range of conditions, most notably ataxias, characterized by the progressive degeneration of these two neuronal populations (Diener & Dichgans, 1992; Fatemi et al., 2013; Sathyanesan et al., 2019). Given the complexity of these circuits, establishing robust human based models is essential to decipher the underlying mechanisms of cerebellar dysfunction. The advent of human pluripotent stem cells (hPSC) has enabled the generation of tissue-specific models to study brain development and disease in a dish. Two-dimensional (2D) cerebellar systems have been successfully used to investigate both development (Madencioglu et al., 2022) and cerebellar-associated diseases (Wong et al., 2017; Xia et al., 2013). However, these models cannot reach full cellular maturation due to their restricted culture duration.

Recent advances in three-dimensional (3D) cerebellar organoids provide promising alternatives (Atamian, Birtele, Hosseini, Quadrato, & Nguyen, et al., 2024; Nayler et al., 2021). Yet, their characterization lags behind more established systems such as cortical organoids (Kadoshima et al., 2013; Lancaster & Knoblich, 2014), particularly regarding early molecular developmental dynamics. In addition, studies addressing early molecular events in human cerebellar development are still limited. Given the cerebellum’s role in higher-order cognitive functions and its human-specific developmental features, further optimization and deeper analysis of these models are needed (Schmahmann, 2019; P. Zhang et al., 2023). Notably, the cerebellum has undergone significant evolutionary expansion in humans, which may have contributed to the emergence of some human-specific traits (Barton & Venditti, 2014). Consistent with this, human Purkinje neurons display distinct developmental dynamics, including an increased production of early-born subtypes, as compared to other species (Sepp et al., 2024).

Here, we generated human pluripotent stem cell–derived cerebellar organoids (CRBOs) and comprehensively characterized their transcriptomic profiles over time, revealing the emergence, diversity, and maturation pace of cerebellar neuronal subtypes. We followed the birth and maturation of glutamatergic and GABAergic neurons to delineate the temporal dynamics of human cerebellar development. Interestingly, our data highlights upstream regulatory factors controlling the pace of cerebellar biogenesis, suggesting important molecular mechanisms that may be altered in cerebellar developmental pathologies. Along this line, we found that cerebellar organoids express genes related to cerebellar pathologies such as ataxia, revealing their potential to study cerebellar diseases *in vitro*.

## Results

### Generation of Cerebellar brain organoids from human PSC

We generated human cerebellum organoids (CRBOs) from H9 cells following a published protocol with minor modifications (Muguruma et al., 2015). Briefly, we introduced slight changes to the culture medium composition and a three-stage static-orbital shaker-bioreactor sequence to increase the growth, survival and development of the organoids (Fig. 1A). CRBOs growth was monitored *in vitro* and showed an increase upon time, as expected (Fig. 1B). To analyze brain regional identity, we compared CRBOs and cortical organoids (COs) for the expression of the early rostral marker OTX2. This analysis of CRBOs at an intermediate stage of 35 days showed almost complete absence of the OTX2 rostral marker among SOX2+ neural progenitors (Fig. 1C-F), in agreement with their caudal identity. In contrast, rostral COs showed a high number of double positive OTX2+ SOX2+ progenitor cells, as expected (Fig. 1G-I). We next compared the expression of rostral forebrain and telencephalic markers such as *TBR1, FOXG1* and *PAX6* and caudal midbrain-hindbrain markers such as *PAX2* and *GAD67* at the transcriptomic level between CRBOs and COs. This analysis revealed that whereas COs showed high expression of rostral regional markers, CRBOs had little or no detectable expression of rostral genes (Fig. 1C,P). On the contrary, CRBOs showed an enrichment of *PAX2* and *GAD67* genes, highlighting their caudal distinctive identity (Fig. 1C,Q). Cerebellar development generates a high diversity of glutamatergic and GABAergic neurons from distinct progenitor populations. SOX2 is a pan-neuronal progenitor marker, whereas PAX6 is a key regulator of neuroectoderm specification and cortical development (Thakurela et al., 2016), also present in cerebellar glutamatergic progenitor and mature neuronal progenies (Muguruma et al., 2015). To assess the presence of cerebellar progenitors in CRBOs, we quantified SOX2+ and PAX6+ cells by immunofluorescence and compared them with corresponding COs at day 35. CRBOs showed a similar proportion of SOX2+ neural progenitors, but a lower percentage of PAX6+ glutamatergic cells (8.25 ± 2.5 %) compared to COs (33.78 ± 4.78 %) (Fig.1J-O,R), consistent with their distinct regional identity. Next, we tested if inclusion of CRBOs and COs within Matrigel scaffolds have an impact on ventricular zone (VZ)-like regions or on the number of SOX2+ and PAX6+ progenitor cells. While Cos cultured in Matrigel showed an increase in VZ surface compared to control conditions, the VZ of CRBOs remained largely unaffected (Fig. S1A-J). Furthermore, no differences in the number of SOX2+; PAX6+ or double positive PAX6+ SOX2+ progenitors were observed with or without Matrigel in both COs or CRBOs (Fig. S1K-L), suggesting that the presence of a scaffold does not have an impact on VZ-like regions, nor on progenitor identity in CRBOs. However, we detected a small increase in the number of KI67+ proliferative cells in CRBOs embedded in Matrigel as compared to controls, suggesting a potential increase in progenitor proliferation in the presence of Matrigel. This difference was not observed when analyzing KI67+ cells within VZ CRBO regions (Fig. S1M-O).

**Figure 1:**
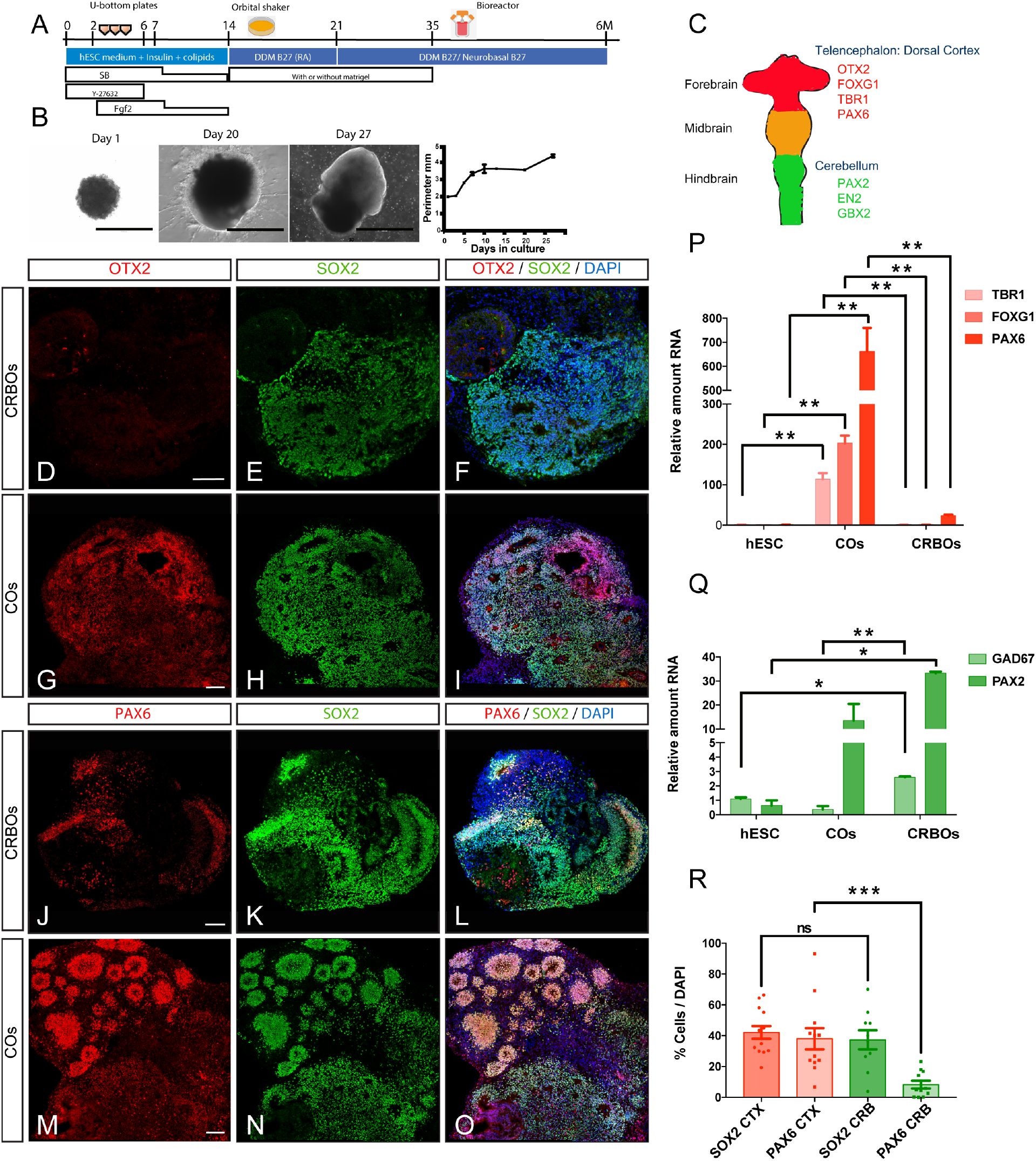
CRBOs show absence of rostral marker expression but presence of glutamatergic progenitors *in vitro*. (A) Scheme of the protocol for the generation of cerebellar organoids (CRBOs). (B) Bright field pictures of brain organoids in culture at days 1, 20, 27 (left). Quantification of brain organoid perimeter (mm) following a timeline in culture (right). Data are shown as Mean ± SEM, (n=3-4). (C) Cartoon depicting the neural tube representation from rostral forebrain (containing the telencephalon: dorsal cortex), to midbrain and hindbrain (containing the cerebellum). (D-O) Immunofluorescence images showing the expression of the rostral progenitor marker OTX2 (in red, D-I), the glutamatergic progenitor marker PAX6 (in red, J-O), and the neuronal progenitor marker SOX2 (green) in CRBOs and COs at day 35. Counterstaining was performed with DAPI (blue). (P-Q) qRT-PCR analysis of the expression of the telencephalic-specific gene FOXG1, and the glutamatergic progenitor genes TBR1, PAX6 (P) and the caudal cerebellar-specific genes GAD67 and PAX2 (Q), in day 0 cells, COs at day 14 and CRBOs at day 14 *in vitro*. Data are shown as relative amount of RNA compared to the values of hESC as value 1 ± SEM (fold change) (n=2). One-way ANOVA with Tukey post-tests, *p<0.05; **p<0.01. (R) Quantification of the percentage of SOX2 and PAX6 cells over DAPI in CRBOs and COs at day 35. Data are represented as mean percentages ± SEM (SOX2: CRBOs 3 Pilots n=12; COs 2 Pilots n=13; PAX6: CRBOs 3 Pilots n=11; COs 3 Pilots n=10). T-test ns= non-significant; ***p<0.001. Scale bars represent 1000 μm (B); and 100 μm (D-O).

Next, we questioned whether 2D and 3D cerebellar differentiation protocols could lead to different outcomes in regional identity. We cultured and differentiated human cerebellar cells in a 2D fashion using similar temporal and culture conditions as for 3D CRBOs (Fig. S2A). Both 2D and 3D differentiation paradigms generated neural progenitors and neurons (Fig. S2B). However, 3D CRBOs showed higher expression of the cerebellar markers *PAX2* and *EN2*, reduced expression of the glutamatergic markers *TBR1, PAX6* and *NEUROD1* as well as absence of the telencephalic marker *FOXG1*, as compared to cells in 2D culture conditions (Fig. S2C-D), suggesting that 3D conditions favor a caudal-cerebellar identity over 2D differentiation paradigms.

These results demonstrate that CRBOs cultured in 3D express key cerebellar genes and lack rostral telencephalic patterning, providing a useful *in vitro* model of the human cerebellum.

### Time-dependent expression of progenitor and neuronal markers in human CRBOs follows the *in vivo* developmental pattern

We analyzed the temporal acquisition of cerebellar regional identities by transcriptomic analysis of CRBOs. We found that CRBOs acquire an early expression of PAX2, EN2 and GBX2 caudal cerebellar regional markers (Lowenstein et al., 2023; Schwarz et al., 1997; Waters & Lewandoski, 2006; Zec et al., 1997) from early time points (14 days) up to 4 weeks in culture (Figure S3A). Moreover, we found a temporal increase in the expression of the progenitor marker SOX1 (Alcock et al., 2009; Pevny & Placzek, 2005) and the neuronal markers NCAM and MAP2 (Johnson & Jope, 1992), suggesting a first peak in neuronal progenitor generation, followed by increased neuronal differentiation (Fig. S3B). Next, we analyzed CRBOs at day 0, day 12-13, day 21-27, and day 35 in culture by bulk RNA sequencing. Principal component analysis (PCA) showed separation of samples according to the number of days in culture (Fig. 2A). Comparative expression analysis revealed a dynamic regulation of genes related to caudal progenitors such as *GBX2, PAX2, EN2, RL* and external granular zone cell precursors (EGZ-GCP) including *ATOH1* (Chang et al., 2019; Lopes et al., 2006; Prasad Poudel et al., 2022), GABAergic progenitors (Lowenstein et al., 2023) such as *PTF1A* and *KIRREL2*, early Purkinje cells (PC) including *OLIG2* and SKOR2 (Ju et al., 2016; Nakatani et al., 2014; Seto et al., 2014); Bergmann glia (He et al., 2026; Hercher et al., 2025; Koirala & Corfas, 2010); and oligodendrocyte precursor cells (OPCs) (Arnett et al., 2004; Stolt et al., 2003), which peaked at intermediate stages (21-27 days) before declining at day 35 (Figure 2B, Fig. S3C-D,F). In parallel, broad markers of neuronal subtypes; DCN neurons such as *LHX2* (Kebschul et al., 2020); mature granule cells (GC) like *PAX6* (Yeung et al., 2016); mature Purkinje cells such as *CALB1* (Lowenstein et al., 2023); oligodendrocyte; and astrocyte cell markers showed a time-dependent increase, consistent with their progressive maturation (Boyles et al., 1985; Eng et al., 2000) (Fig. 2B, Fig. S3E,G-H). In order to confirm that the changes that occur in the *in vitro* CRBOs parallel those that happen in the human brain, we compared the dataset with the BRAINSPAN atlas (https://www.brainspan.org/) (Miller et al., 2014). This comparison showed a similar temporal pattern of expression of immature neuronal progenitors and neuronal progenies between CRBOs and human embryonic cerebellar tissue (Fig. 2B-C, Fig. S3C-H).

**Figure 2:**
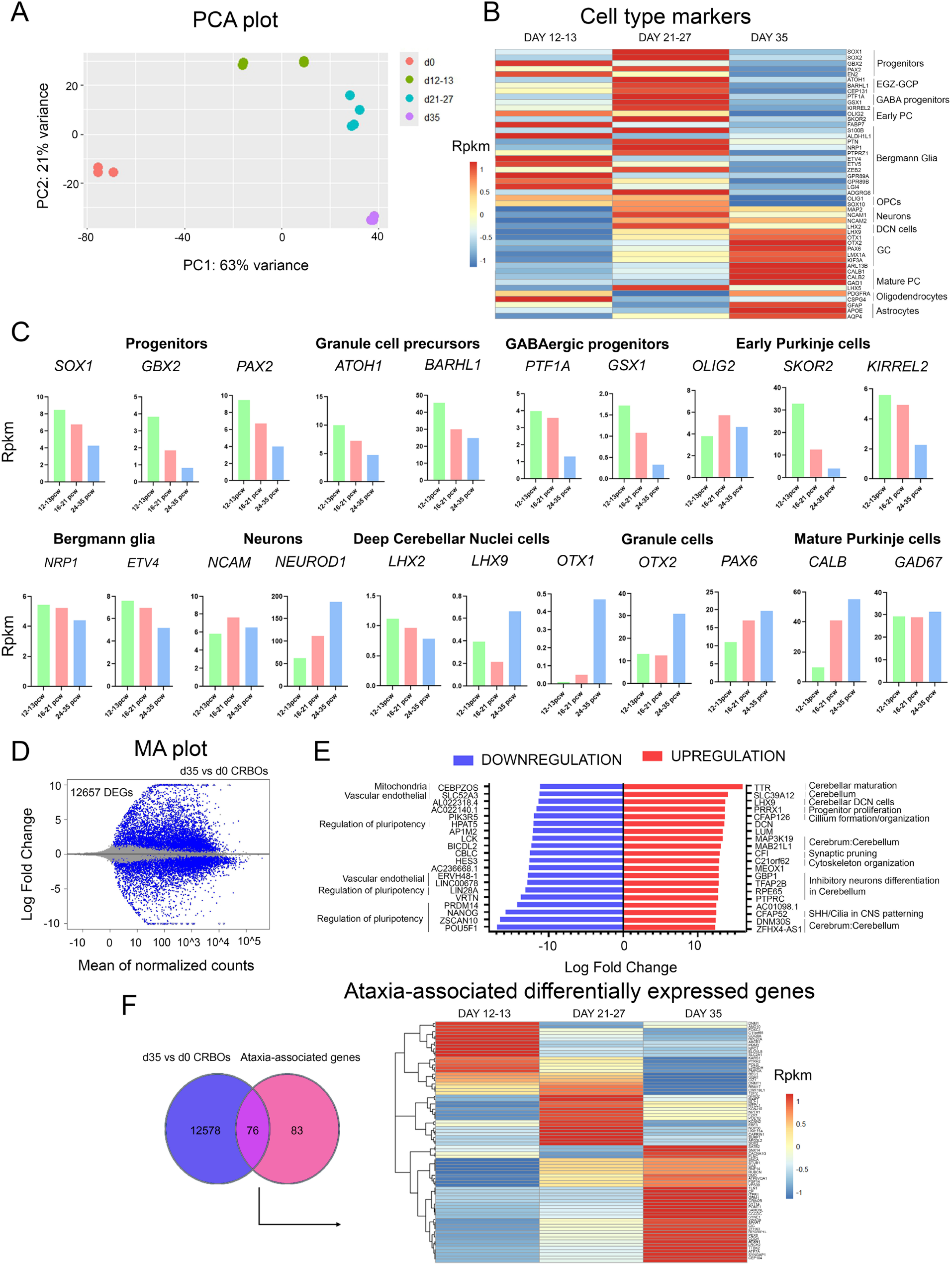
Transcriptomic profile of *in vitro* CRBOs confirms their cerebellar identity. (A) PCA showing the different dynamics of the different samples sequenced by bulk RNA-seq. The x-axis represents the principal component 1 (PC1) that separates with 63% of the variance the day0 samples in the left and the day 12-13, day 21-27 and day 35 in the right. The y-axis represents the PC2 with 21% of variance that separates between day35 and day 12-13, and day 21-27. (B) Heatmap showing rpkm levels from bulk RNA-seq performed in day 12-13, day 21-27 and day35 *in vitro* CRBOs samples. EGZ-GCP= External Germinal Zone of Granule Cells Precursors, PC= Purkinje Cells, OPCs=Oligodendrocyte Precursor Cells, DCN=Deep Cerebellar Nuclei, GC=Granule Cells. Rpkm = Read per kilobase million. (C) Expression levels as Rpkm obtained from the BRAINSPAN database. pcw: post conception weeks. mos: months. (D) MA plot showing DEGs in the CRBOs at 35 days compared to hESC *in vitro*. Each blue dot represents one DEG defined by FDR < 0.05. The x-axis represents the mean of normalized counts and the y-axis the log2-fold change. d35=35 days *in vitro* CRBOs. (E) Back-to-back bar plot showing top20 downregulated (blue) and upregulated (red) DEGs when comparing CRBOs at 35 days *in vitro* with hESC. Cell-type enrichment analysis was done using these genes as input for DESCARTES database and filtering by Central Nervous System. (F) Venny plot (left) and heatmap (right) showing the 76 shared genes among DEGs obtained from day35 versus hESC and ataxia-associated genes downloaded from DISGENET. ATXN1 highlighted in Bold.

These results show that CRBOs in culture undergo a first wave of progenitor proliferation followed by cell cycle exit of the progenitors to generate most neuronal and glia progenies, similarly to *in vivo* cerebellar development.

### CRBOs show developmentally regulated changes in cytoskeleton, neuronal activity, plasticity and SHH-associated cilia processes

Next, we performed a differential expression analysis comparing day 0 and day 35 CRBOs. This analysis revealed 12657 differentially expressed genes (DEGs) (Figure 2D, Supplementary Table 1). Among the top 20 upregulated genes we found categories such as cerebellar identity (Figure 2E, Supplementary Table 2); markers of excitatory DCN cerebellar cells like *LHX9;* and inhibitory neuronal markers (Figure 2E) *(Jin et al*., *2015)*. We also found upregulation of genes involved in progenitor proliferation, synaptic pruning (Gomez-Arboledas et al., 2021; J. Wang et al., 2018), cytoskeleton and cilium formation (Figure 2E). Top 20 downregulated genes included mitochondria and regulation of pluripotency related genes, suggesting a higher differentiation stage at day 35 in CRBOs (Figure 2E).

To evaluate the utility of CRBOs for modeling cerebellar-associated diseases, we assessed the overlap of ataxia-associated genes from the DisGeNET database (https://disgenet.com/) (Piñero et al., 2015) with the DEGs at day 35 *vs* day 0 CRBOs (Supplementary Table 3). Interestingly, over 50% of known ataxia genes (76/159) were represented among the CRBO DEGs, including the key inherited ataxia gene *Ataxin 1* (*ATXN1*) (Fig. 2F, Supplementary Table 1).

Next, we assessed the progression of the cerebellar differentiation by performing differential expression analysis between early, day 12–13, and the intermediate time point day 35 (Fig. 3A, Supplementary Table 4). Enrichment analysis and Ingenuity Pathway Analysis (IPA) (Krämer et al., 2014) for upregulated DEGs revealed lysosome, axonal guidance, focal adhesion, SHH, integrin changes and extracellular matrix (ECM)-receptor interaction among the top enriched pathways (Fig. 3B-E, Supplementary Table 5-8). All these pathways were associated with a higher presence of neurons and their synaptic network at day 35 in CRBOs. Of those, cilium assembly and organization, and cilium-dependent motility, were among the most significant enriched pathways (Fig. 3C, Supplementary Table 6), in line with the developmental role of the primary cilium in EGZ-GCPs, which concentrates components of the SHH signaling. SHH is secreted during cerebellar development by PCs to promote the proliferation of EGZ-GCPs (Spassky et al., 2008). In agreement with this, we also found SHH among the top15 most enriched pathways in day 35 CRBOs when compared to day 12-13 (Fig. 3B, Supplementary Table 5). Additionally, we also found Reelin signaling among upregulated pathways (Supplementary Table 5), consistent with its role in the migration and organization of PCs and GCs within the IGL. Interestingly, we observed the enrichment of potassium channels and CREB signaling (Fig. 3E, Supplementary Table 8), which are associated with neuronal plasticity and activity (Dworkin & Mantamadiotis, 2010; Sakamoto et al., 2011), and regulation of hPSC pluripotency.

**Figure 3:**
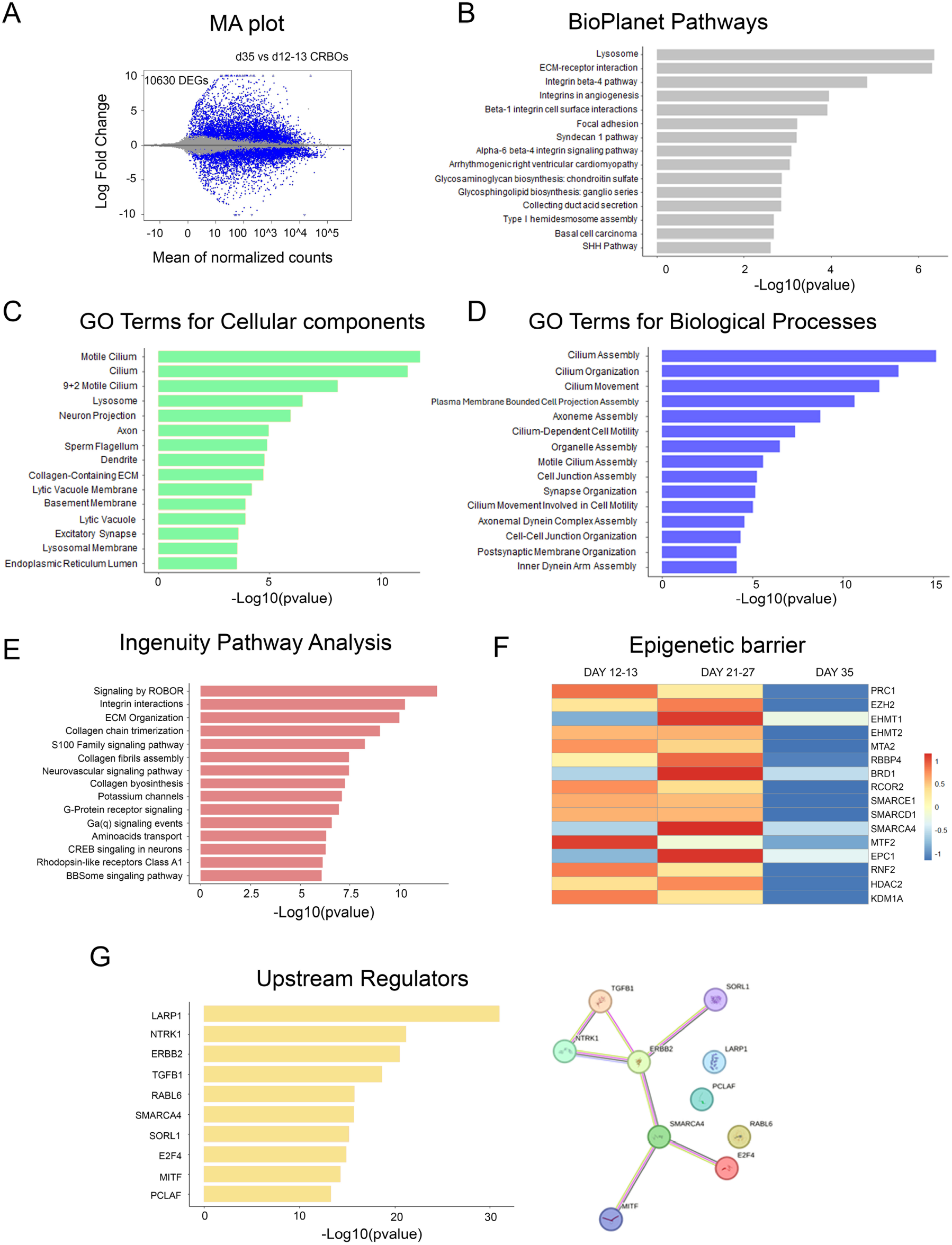
Time-dependent transcriptomic profile of *in vitro* CRBOs show upregulation of neuronal and SHH-associated cilia processes, and downregulation of epigenetic factors. (A) MA plot showing DEGs in the CRBOs at 35 days compared to12-13 days *in vitro*. Each blue dot represents one DEG defined by FDR < 0.05. The x-axis represents the mean of normalized counts and the y-axis the log2-fold change. D12-13=12 and 13 days *in vitro* CRBOs. d35=35 days *in vitro* CRBOs. (B-D) Pathway enrichment analysis for the 5333 upregulated DEGs in day 35 *in vitro* CRBOs compared with day 12-13 *in vitro* CRBOs using Enrichr. The top 15 results from (B) BioPlanet pathways, (C) GO terms for cellular components, and (D) GO terms for biological processes based on the significance of enrichment are shown. (E) Pathway enrichment analysis over all DEGs (after applying a cut off of absolute fold change > 1) in day 35 compared with day 12-13 *in vitro* CRBOs using IPA and selecting the upregulated ones based on positive Z score. (F) Heatmap showing rpkm levels from bulk RNA-seq performed in day 12-13, day 21-27 and day35 *in vitro* CRBOs samples. Epigenetic Factors associated with regulation of human neuronal maturation appear downregulated at later time points. (G) Left: Top 10 upstream regulators based on the significance of enrichment predicted by IPA according to the DEGs (after applying a cut off of absolute fold change > 1) from day 35 versus day 12-13 *in vitro* CRBOs. Right: Connections observed between the top 10 upstream regulators predicted by IPA. SORL1, ERBB2, TGFB1, NTRK1, SMARCA4, E2F4 and MTF are directly or indirectly connected through different pathways, while there is no evidence of connection with LARP1, PCLAF and RABL6.

These results overall show that 35 days CRBOs have increased maturation stage compared to earlier time points and a transcriptomic gene expression pattern consistent with cerebellar progenies.

### CRBOs show loss of epigenetic modulators during differentiation *in vitro*

Differential expression analysis comparing days 12–13 and day 35 *in vitro* CRBOs further revealed a downregulation of genes related to translation, gene expression, and cell cycle regulation (Figure S4A-E, Supplementary Table 5-7). In addition, multiple mitochondria-associated pathways were among the most significantly downregulated categories (Figure S4C, D Supplementary Table 6-8), indicating substantial molecular and cellular remodeling during cerebellar differentiation.

These findings are consistent with a tight regulation of cell cycle gene expression during development, where its activity decreases as cells exit the pluripotent state and undergo differentiation. This temporal upregulation of maturation genes and downregulation of pluripotency and cell cycle related genes may reflect a reduced expression of epigenetic regulators that set the timing of human cerebellar maturation (Aldridge & West, 2024; Ciceri et al., 2024; Nowakowski et al., 2017). Such factors constitute an epigenetic barrier that is progressively released over time, thereby allowing cell and tissue maturation (Ciceri et al., 2024). In line with this observation, we found a time-dependent downregulation of epigenetic regulators associated with this barrier in CRBOs such as *Protein Regulator of Cytokinesis 1 (PRC1), Metastasis Associated 1 Family Member 2 (MTA2), Metal Response Element Binding Transcription Factor 2 (MTF2), and Lysine Demethylase 1A (KDM1A)*, among others (Fig. 3F), further supporting a molecular progression toward a more advanced maturation state.

These data suggest that differentiating CRBOs generate progressively more mature progenies through downregulation of epigenetic factors.

### Predictive upstream analysis reveals the role of proteins associated with mitochondria, cell cycle and brain development

To identify the upstream drivers of the transcriptomic changes observed at intermediate stages (day 35) versus early stages (Days 12 and 13) of CRBO maturation, we performed an upstream regulator prediction using IPA on DEGs. Top 5 predicted upstream regulators included *La Ribonucleoprotein 1*, Translational Regulator (*LARP1*), a well-known regulator of translation (Philippe et al., 2020) (Fig. 3G, Supplementary Table 9), in line with pathway enrichment results, *Neurotrophic Receptor Tyrosine Kinase 1* (*NTRK1*), *Erb-B2 Receptor Tyrosine Kinase 2* (*ERBB2*) and *Transforming Growth Factor Beta 1* (*TGFB1*) (Fig. 3G). In agreement, tyrosine kinases play an essential role during cerebellum development as regulators of neuronal migration and cerebellar layer formation (Boxall et al., 1996; Canepari & Ogden, 2003; Qiu et al., 2010). Indeed, NTRK1 is associated with higher neural activity (Shmakova et al., 2021), while ERBB receptors and TGFB1 are critical for nervous system development (Birchmeier, 2009; Dalvand et al., 2022). TGFB1 regulates early cerebellar development through modulation of cell adhesion and autophagy (Dalvand et al., 2022). Protein–protein interaction network analysis using STRING (von Mering et al., 2005) revealed a close connectivity between *NTRK1, ERBB2* and *TGFB1* (Fig. 3G). Additional top regulators interconnected in the STRING network included *SWI/SNF Related, Matrix Associated, Actin Dependent Regulator of Chromatin, Subfamily A, Member 4* (SMARCA4), *E2F Transcription Factor 4* (*E2F4*), and *Microphthalmia-Associated Transcription Factor* (*MITF*). Of those, *E2F4* is involved in cell cycle regulation and brain development, whereas *MITF* contributes to cellular redox homeostasis (Lichtlen et al., 1999; Rutherford & Bird, 2004), suggesting that redox balance may be critical for the transition from progenitors to immature neurons. Interestingly, *SMARCA4* is a chromatin remodeler required for synapse development (Z. Zhang et al., 2016) and modulator of the SHH pathway (Shi et al., 2016), that was also identified as an epigenetic barrier factor (Fig 3F).

These data suggest that generation of CRBOs *in vitro* recapitulates the expression of upstream pathways controlling neural differentiation, cerebellum development, cell cycle regulation, redox balance and epigenetic regulation.

### CRBOs showed temporal biogenesis of Glutamatergic cerebellar neurons

Next, we characterized the glutamatergic populations in CRBOs following long-term culture (up to 6 months to ensure the development of more mature neuronal progenies). Glutamatergic neuronal progenies are differentiated from glutamatergic progenitors located within the Rhombic lip (RL) of the cerebellum. Cerebellar glutamatergic neuronal populations include cerebellar nuclei neurons within the deep cerebellar nuclei (DCN); unipolar brush cells (UBCs); and granule cells (GC). DCN neurons are the first glutamatergic neurons born in the cerebellum, followed by UBCs and GCs. At the molecular level, distinct glutamatergic progenies are identified based on the expression of transcription factors such as *TBR1* and *LHX2* which mark DCNs (Fink et al., 2006; Kebschul et al., 2020; Yeung et al., 2016), or PAX6 which is enriched in GCs (Yeung et al., 2016). Our results show that LHX2+ DCN neurons are detected as early as day 27-35 and are present in late d49-63 CRBOs (Fig. 4A-P). Interestingly, we detected an increase in LHX2+ DCN glutamatergic neurons in day 49-63 CRBOs (18.02 ^±^ 3.62 % of total DAPI) compared to day 27-35 (6.69 ± 1.96 %) (Fig. 4Q), suggesting a progressive differentiation of DCN neurons upon time in culture. In agreement with this, we detected the colocalization of LHX2 and TBR1 TFs at day 63 in CRBOs, confirming their DCN glutamatergic identity (Fig. 4R-T). In addition, we identified the presence of numerous postmitotic PAX6+; SOX2-; MAP2+ neurons at day 49-63 and in long-term 5-6M CRBOs (Fig. 4U-X), corresponding to cells with GC identity.

**Figure 4:**
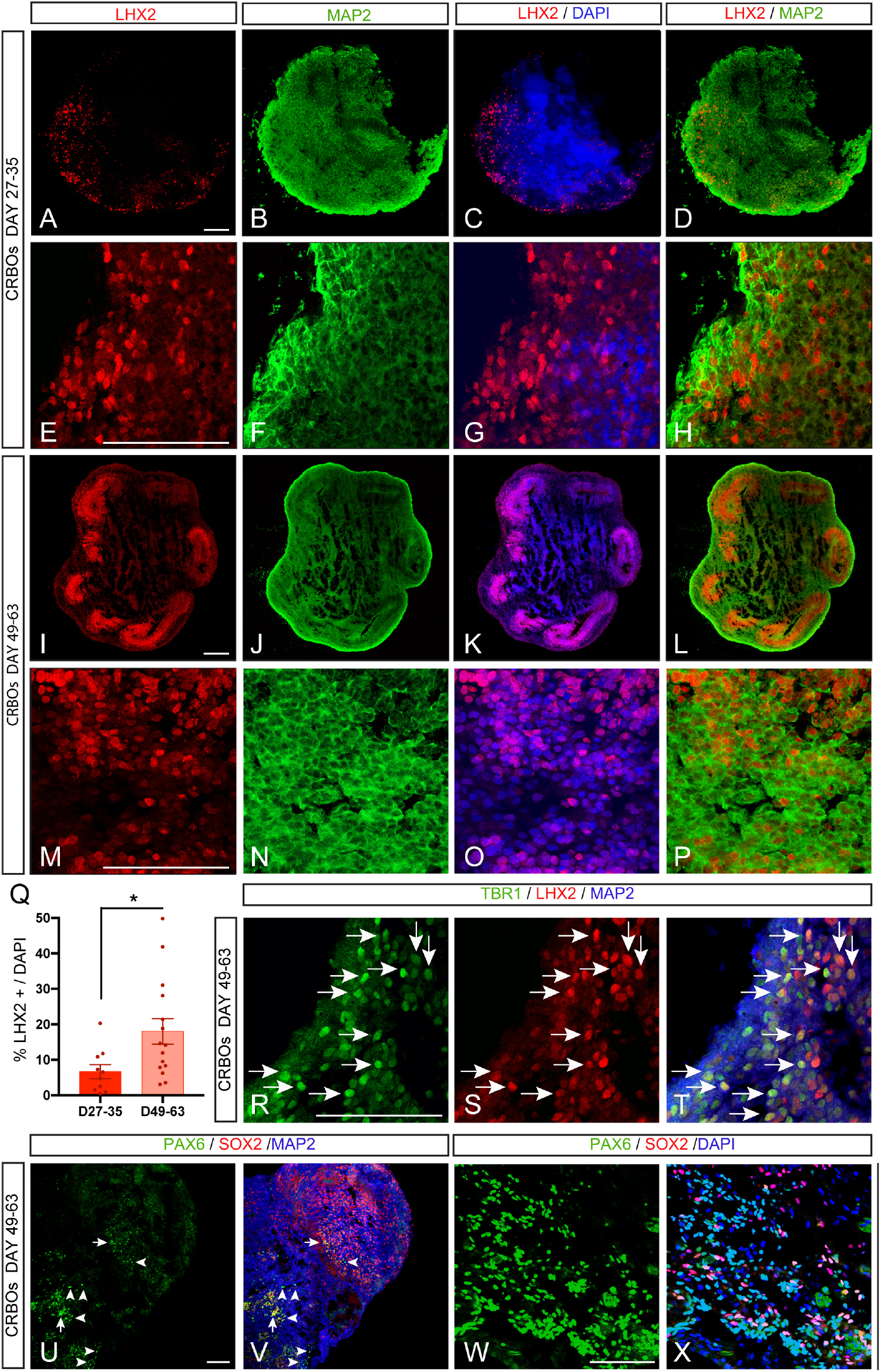
Long-term CRBOs recapitulate the presence of glutamatergic neuronal populations such as DCN and GC. (A-P) Tile scan (A-D,I-L) and high magnification (E-H,M-P) immunofluorescence images showing the expression of the glutamatergic DCN marker LHX2 (in red), and the neuronal marker MAP2 (in green) in day 27-35 CRBOs (A-H) and day 49-63 CRBOs (I-P). Counterstaining was done with DAPI (blue). (Q) Quantification of the proportion of LHX2+ cells among DAPI in CRBOs at day 27-35 and day 63. Data are represented as mean percentages ± SEM (CRBOs d27-35 3 Pilots n=10; d49-63 3 Pilots n=15). T-test * p<0.05. (R-T) Immunofluorescence images showing the co-localization of TBR1 (in green) and LHX2 (in red) in DCN glutamatergic neurons in long-term CRBOs (white arrows). (U-X) Tile scan (U-V) and single (W-X) immunofluorescence images showing the presence of PAX6+ SOX2+ glutamatergic progenitors (white arrows) and PAX6+ SOX2-glutamatergic GC (white arrowheads) in day 49-63 CRBOs (U-V) and in long-term 5-6M CRBOs (W-X). Scale bars represent 100 μm.

These data suggest that CRBOs recapitulate the generation of Glutamatergic populations corresponding mostly to DCN and GC identities from the cerebellum.

### CRBOs showed an increase in GABAergic cerebellar neurons upon time

Finally, we sought to identify GABAergic cerebellar progenies present in CRBOs upon time in culture. Cerebellar GABAergic neurons are differentiated from GABAergic progenitors residing in VZ regions *in vivo (Lowenstein et al*., *2023)*. Cerebellar GABAergic neurons include small GABA DCN; Purkinje cells, and small inhibitory neurons (stellate cells and Golgi neurons). First, we analyzed by immunofluorescence the presence of OLIG2+; SOX2+ PC progenitor cells inside VZ-like regions and early postmitotic OLIG2+; SOX2-PC outside of VZ-like regions. This analysis revealed the presence of both PC populations at day 35 CRBOs (Fig. 5A-C). In order to identify more mature PC populations, we looked at the temporal expression of CALB and GAD67 in CRBOs. We detected the presence of numerous GAD67+ interneurons and CALB+ PC mature neurons in CRBOs at various time points in culture, in contrast with the low presence or absence of staining in rostral COs, as expected (Fig. 5D-O, Fig. S3G-N). These experiments showed the presence of GAD67+; CALB-; MAP2+ inhibitory neurons in CRBOs next to the population of GAD67+; CALB+; MAP2+ mature PC in CRBOs (Fig. 5D-L). Next, we compared the relative proportion of CALB+ mature PC upon time in culture. We found a relative increase in the organoid area covered by CALB+ staining in day 49-63 CRBOs (4.59 ± 0.35 %) compared to early d13-14 CRBOs (2.39 ± 0.51 %) *in vitro*, suggesting an increased differentiation of PC neurons with time in culture (Fig. 5M). Lastly, we identified the presence of PC mature neurons in long-term 5-6M CRBOs (Fig. 5N-O).

**Figure 5:**
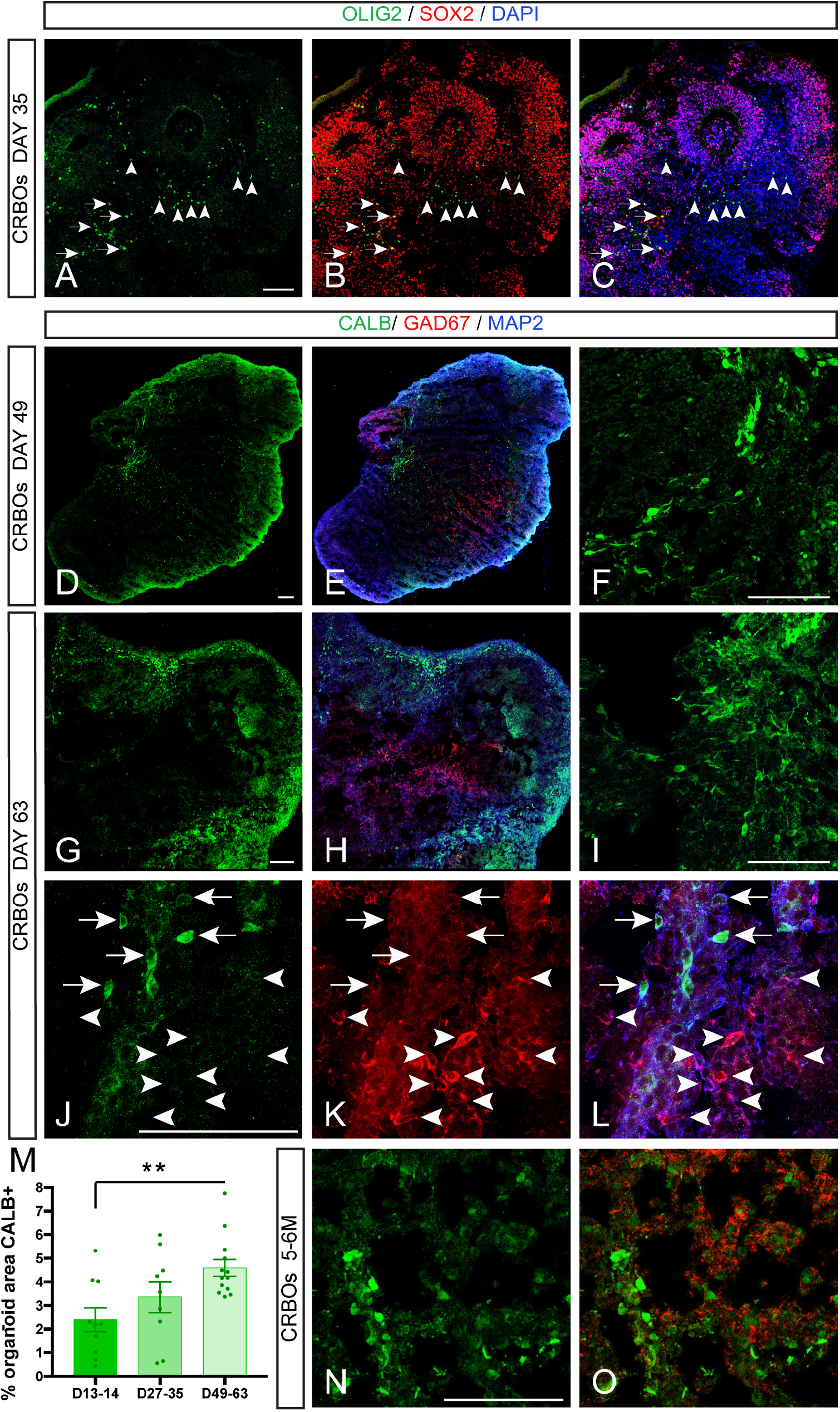
CRBOs show the presence of GABAergic neuronal progenies such as PC and other interneurons. (A-C) Immunofluorescence tile scan images showing the presence of OLIG2+ SOX2+ PC progenitors (white arrows) and early postmitotic OLIG2+ SOX2-PC cells (white arrowheads) in day 35 CRBOs *in vitro*. (D-L, N-O) Tile scan (D-E, G-H) and single (F, I, J-L) immunofluorescence images showing the presence of mature CALB+ MAP2+ PC (in green) and GAD67+ CALB-MAP2+ interneurons (in red) in day 49-63 CRBOs (D-F), day 63 CRBOs (G-L) and long-term 5-6M CRBOs (N-O). White arrows mark CALB+ GAD67+ MAP2+ mature PCs, whereas white arrowheads highlight CALB-GAD67+ MAP2+ GABAergic interneurons in day 49-63 CRBOs (J-L). (M) Quantification of the CALB+ organoid area in CRBOs at 13-14; 27-35 and 49-63 days. Data are represented as mean percentages ± SEM (CRBOs d14 3 Pilots n=10; d27-35 3 Pilots n=10; d49-63 3 Pilots n=13). One-way ANOVA ** p<0.01. Scale bars represent 100 μm.

These results show that long-term CRBOs can recapitulate to some extent in vivo cerebellar development with the presence of various GABAergic neuronal subtypes, including non-PC inhibitory neurons and mature PC.

## Discussion

In this study, we generated cerebellar organoids from human pluripotent stem cells and characterized their molecular and cellular changes upon time *in vitro*. Through combined transcriptomic and immunofluorescence analyses, we demonstrate that CRBOs acquire an early caudal identity characterized by the absence of rostral markers and the enrichment of midbrain-hindbrain-specific genes, followed by acquisition of cerebellar-specific regional identity. Caudal specification of CRBOs was further validated through comparison with cortical organoids (COs), which exhibit rostral identity.

Temporal transcriptomic profiling of CRBOs across developmental stages revealed a progressive transition from progenitor populations to differentiated neuronal and glial lineages. Early enrichment of cerebellar progenitor markers, including those associated with Bergmann glia, glutamatergic and GABAergic lineages, was followed by increased expression of markers corresponding to mature neuronal populations, such as DCN, GCs, and PCs. Notably, this temporal pattern of expression paralleled that from prenatal human cerebellar tissue, highlighting the ability of CRBOs to recapitulate key features of early human cerebellar development. In this work, CRBOs were generated following a directed patterning method with TGFβ inhibition to induce neuroectoderm differentiation, combined with FGF2 and insulin to induce caudal identity. This paradigm has been used previously to derive cerebellar organoids, however, a co-culture with mouse-derived external granular layer cells was required to generate PCs (Ishida et al., 2016; Muguruma et al., 2015). In contrast, our system employs a three-stage static-orbital shaker-bioreactor sequence to improve survival, growth and long-term culture of CRBOs and its neuronal progenies such as PC progenitors and postmitotic PCs, without a 2D intermediate step. While we and others (Nayler et al., 2021) show that the combination of FGF2 and insulin promote caudal cerebellar specification, a recent report described forebrain identity acquisition following this paradigm (Atamian, Birtele, Hosseini, & Quadrato, 2024). Instead, the authors used a combination of dual SMAD inhibition, WNT activation and FGF8 to generate cerebellar organoids. These findings suggest that multiple morphogen combinations can converge to generate a cerebellar identity *in vitro*.

Although prior single-cell RNA-seq studies demonstrated the presence of diverse cerebellar cell types in CRBOs (Nayler et al., 2021), a description of its temporal dynamics was largely missing. Here, we provide a detailed temporal framework of the generation of cerebellar progenies *in vitro*, showing an early acquisition of caudal cerebellar identity at 2 weeks, followed by peak generation of glutamatergic and GABAergic progenitors at 4 weeks, and subsequent differentiation into postmitotic neuronal and glial populations by 5 weeks *in vitro*. Differentiation of organoids that mimic other brain regions, such as the cortex, have previously used Matrigel to support growth of COs. To understand if Matrigel may be necessary to support CRBO growth or cell identity, we differentiated CRBOs and COs in the presence or absence of a Matrigel scaffold. We found that whereas COs showed larger VZ-like regions when embedded in Matrigel, CRBO VZ area was largely unaffected. Moreover, we found no changes in progenitor cell identity between both conditions whether in COs or CRBOs, suggesting a negligible or no contribution of a scaffold into cerebellar differentiation. Our results are in contrast with a previous study showing that the encapsulation of cerebellar organoids led to increased generation of glutamatergic neurons, albeit with higher heterogeneity than controls (Nayler et al., 2021). The discrepancies between our findings and the previous study may stem from our specific focus on progenitor identities, which remained unaffected. Additionally, the high degree of heterogeneity introduced by the Matrigel scaffold likely contributed to the differing outcomes.

At the molecular level, the differentiation of CRBOs involved an upregulation of cytoskeletal remodeling, extracellular matrix interactions, axon guidance, lysosomal function, and neuronal activity pathways, reflecting progressive structural and functional maturation. Notably, cilium-related processes and Sonic Hedgehog (SHH) signaling were strongly enriched, supporting active granule cell progenitor proliferation. In parallel, Reelin signaling was also upregulated, consistent with its role in neuronal migration and laminar organization during cerebellar development. Conversely, genes associated with translation, RNA processing, and cell cycle regulation were downregulated over time, in line with the transition from proliferative progenitors to differentiated cells (Nguyen et al., 2002). We further observed a coordinated downregulation of epigenetic regulators previously implicated in neuronal maturation tempo (Ciceri et al., 2024), supporting the concept of a progressively diminishing “epigenetic barrier” that permits differentiation. In addition, mitochondrial and oxidative phosphorylation pathways were reduced in intermediate *in vitro* stages, suggesting that CRBOs at 35 days remain metabolically immature, reflecting still an early stage of neuronal development. Future studies extending culture duration will be important to assess the transcriptome of later maturation stages.

Finally, we identified predicted upstream regulators of temporal transcriptomic shifts in CRBOs. These include factors involved in translation, cell cycle control, cellular redox maintenance, and SHH pathway components.

Our study demonstrates a time-dependent increase in both glutamatergic and GABAergic neuronal populations within CRBOs over a two-month period. Among glutamatergic progenies, DCN neurons are the earliest-born within the cerebellum, and accordingly, LHX2^+^ neurons were detected in CRBOs as early as day 27–35 and increased in proportion by day 63. The co-expression of LHX2 and TBR1 transcription factors further confirmed the identity of these cells as DCN glutamatergic neurons. In addition, day 63 and long-term 5-6M CRBOs contained numerous postmitotic PAX6^+^ SOX2^−^ MAP2^+^ neurons corresponding to granule cell identity, which represent the most abundant neuronal population of the cerebellum. Together, these findings indicate that CRBOs recapitulate the generation of major cerebellar glutamatergic neuronal populations and suggest that the temporal progression of rhombic lip-derived progenies is preserved in this *in vitro* system. In parallel, CRBOs also generated GABAergic neuronal populations that arise from ventricular zone progenitors during cerebellar development. Indeed, OLIG2^+^; SOX2^+^ Purkinje cell progenitors within ventricular zone-like regions as well as early postmitotic OLIG2^+^; SOX2^−^ cells were detected, indicating the presence of both proliferative and postmitotic Purkinje cell populations. Consistent with progressive neuronal maturation, 2 months old CRBOs showed an increased proportion of CALB^+^ mature Purkinje cells compared to earlier stages. In addition to Purkinje neurons, we also identified GAD67^+^ inhibitory interneurons lacking CALB expression, indicating the presence of non-Purkinje inhibitory neuronal subtypes. Notably, the coexistence of these two populations - GABAergic alongside glutamatergic neurons - underscores the ability of CRBOs to recapitulate key features of cerebellar neuronal diversity and maturation *in vitro*. Together, these results demonstrate that extended culture of CRBOs supports the differentiation of both major cerebellar lineages essential for the establishment of cerebellar neuronal circuitry.

Lastly, CRBO differentiation was accompanied by a time-dependent upregulation of genes associated with cerebellar disorders, including key inherited ataxia genes such as *ATXN1*. Further, these findings suggest that CRBOs could provide a valuable platform for modeling cerebellar-associated disorders and to investigate early molecular events underlying disease development.

In a nutshell, the CRBO model described here, together with its detailed molecular characterization, may provide a valuable platform to investigate the mechanisms underlying neurodevelopmental cerebellar disorders as well as the early molecular events that precede adult-onset cerebellar diseases such as ataxia in a human context. Indeed, brain organoids representing other brain regions, including cortical and midbrain organoids, have already been successfully used to study neurological diseases and developmental disorders (Bubnys & Tsai, 2022; Di Lullo & Kriegstein, 2017; Eichmüller & Knoblich, 2022; Smits et al., 2020). In this context, the establishment and characterization of human cerebellar organoids open new opportunities for modeling cerebellar pathologies and for developing drug-screening approaches aimed at understanding how cerebellar cell populations and their maturation may be affected by therapeutic interventions in the early stages of the disease, ultimately accelerating progress in cerebellar disease research.

## Acknowledgements

This work was supported by the Credit Classique from the University of Liège (Credit Classique 2020 (to I. E.- C.)), the WBI Excellent Grant from the Federation Wallonie Bruxelles (FWB to S.H.), the F.R.S.-FNRS (#CUR 40002797 (to L. N. and I. E.- C.), and PER 40003579 (to L. N., I. E.-C.). L.N. is Research Director of the F.R.S.-F.N.R.S. We thank the GIGA Imaging, GIGA Genomics and GIGA Bioinformatics platforms for their contribution, help and support to this work.

## Disclosure of potential conflict of interests

The authors declare no competing interests.

## Author contributions

I.E-C. designed the experimental plan. S.H., A.B., I.E-C. performed the experiments with the help of N.O., B.C. S.H., A.B., I.E-C. analyzed all the results. S.H., I.E-C. wrote the manuscript with the help of A.B. and the rest of coauthors. I.E- C. and L.N. designed the initial study.

## Materials and Methods

### Differentiation of CRBOs from hPSC

Cerebellum brain organoids were developed following a modified version of an already existing protocol (Muguruma et al., 2015). hESC-H9 cell line (Wicell, WA09, metadata: female, no disease associated) was dissociated using accutase (Stem Cell Technologies: 7922) and 9000 cells were seeded in 96 well U bottom plates (Nunclon sphera: 15396123). Cells were cultured in DMEM-F12 with 20% KO serum (ThermoFisher), NEAA; penicillin/streptavidin and 100µm β-mercaptoethanol for 14 days. Morphogens were added to the medium for 14 days: 10µM SB (SB431542; Sigma: S4317) and 7µg/ml insulin (Sigma: I9278-5ML), chemically defined lipid concentrate (1% Invitrogen). From day 2, 50µg/ml FGF2 (Peprotech: 100-18b) was added. After day 7, the concentration of SB and FGF2 were reduced to 5µM and 25µg/ml, respectively. Rock inhibitor (Y-27632 2HC; Bio-Connect: S1049) was added on day 0 at 20µM and at 10µM from day 2 to day 6. From day 14 to day 28, medium was composed of DMEM-F12, sodium pyruvate (ThermoFisher: 11360-039), NEAA, N2 (ThermoFisher: 17502048), Pen/strep, BSA (ThermoFisher: 9048-46-8), B27 without vitamin A (ThermoFisher: 11500446) and 100µm β-mercaptoethanol. Insulin (7µg/ml) and 1% co-lipids were added to the medium until day 21. On day 23-25, organoids were transferred to 6cm dishes and placed on orbital shakers (75 rpm). From day 28 to day 35, medium was composed of half DMEM-F12 and half Neurobasal (ThermoFisher: 21103-049), supplemented with Na Pyr, NEAA, N2, Glutamax (ThermoFisher: 35050-038), BSA, Pen/strep, B27 without vitamin A and 50µm β-mercaptoethanol.

### RNA sequencing and analysis

High quality RNA samples with a RQN value above 7 were used for sequencing: 3 samples day 0, 2 samples day 12, 2 samples day 13, 2 samples day 21, 2 samples day 27 and 3 samples day 35. mRNA libraries were generated by the GIGA Genomics Platform (www.gigagenomics.uliege.be) using the TruSeq Stranded mRNA Library Prep kit (Illumina) and sequenced using the Illumina Novaseq 6000 machine to generate paired-end 2×150bp reads. Downstream analysis was performed using STAR for mapping to the human reference genome GRCh38 release 103 and Salmon to produce the counts of the reads. Raw counts were processed using dplyr package to generate the reads per kilo base of transcript per million mapped reads (rpkm). Raw counts were processed using DESeq2 R package (v1.36.0) (Love et al., 2014) to perform the differential expression analysis and obtain the differentially expressed genes (DEGs).

### Statistical analysis

All statistical analysis was performed using GraphPad (Prism). All data were tested for normality using the Shapiro-Wilcoxon test. All information concerning the test used and the multiple comparisons test applied can be found in the legend of the corresponding figure.

### Data and code availability

The RNA-seq data for this study have been deposited in the European Nucleotide Archive (ENA) at EMBL-EBI under accession number PRJEB…

RNA-seq analysis codes can be found in the following repository: https://github.com/Espuny-Camacho/CEREBELLUM_ORGANOIDS/

## Supplemental Information

### Supplemental Materials

#### Supplemental Figures

**Figure S1:**
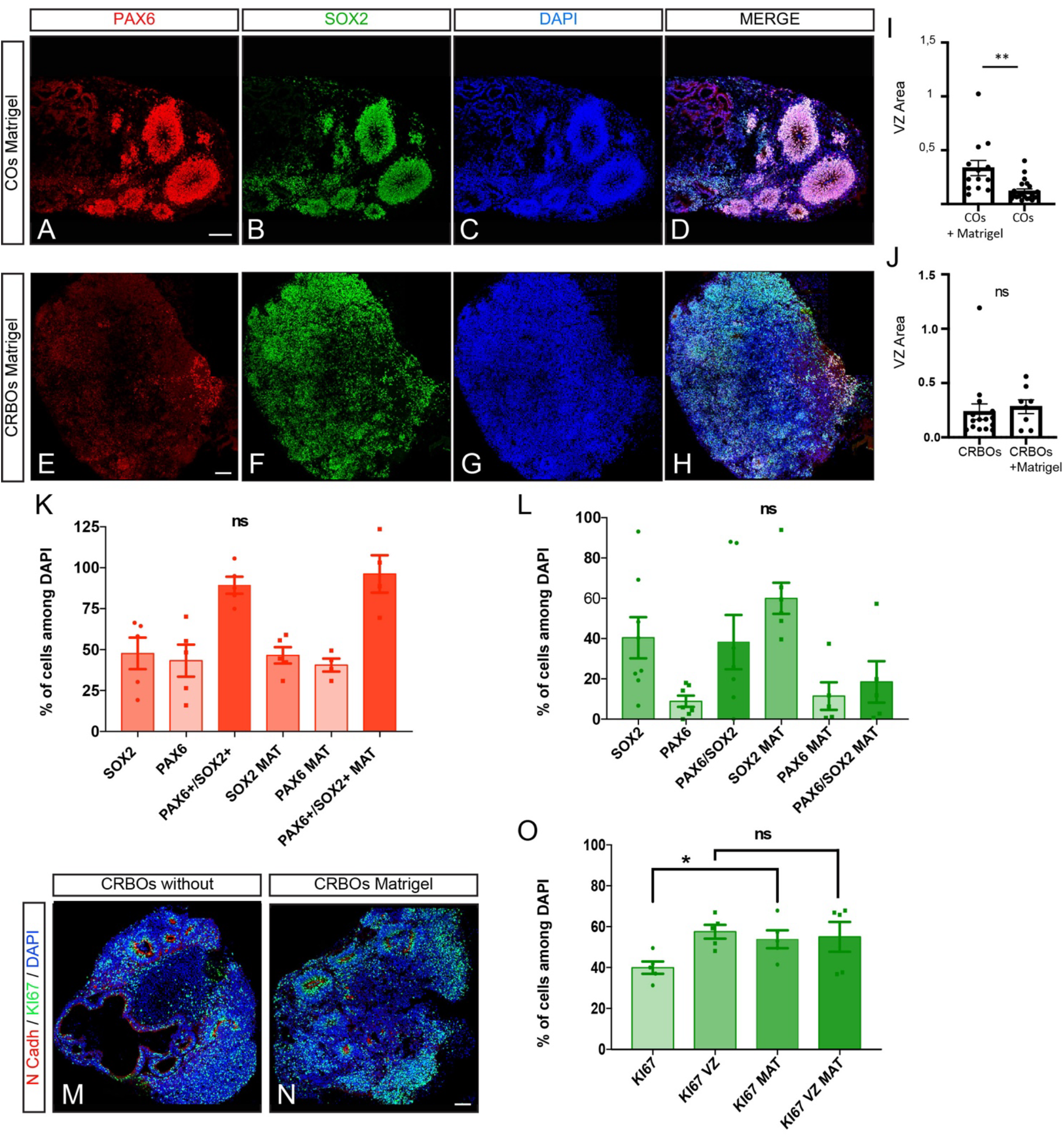
Presence of an organoid scaffold does not change cellular identity in CRBOs or COs *in vitro*. (A-H) Immunofluorescence tile scan images showing the expression of the glutamatergic marker PAX6 (red) and the neuronal progenitor marker SOX2 (green) in COs and CRBOs embedded in matrigel. (I-J) Quantification of the number and area of VZ regions (in μm^2^) in COs (I) and CRBOs (J) in the presence or absence of matrigel. Data are represented as mean ± SEM (COs n=4-5; CRBOs n=5-8). (K-L) Quantification of the percentage of SOX2+; PAX6+; PAX6+/SOX2+ in COs (K) and CRBOs (L) in absence or presence of Matrigel. (COs n=4-5; CRBOs n=5-8). Data are represented as mean ± SEM. T-test ns= non-significant. (M-N) Immunofluorescence images showing the expression of the apical marker N-CADHERIN (red) and the proliferative marker KI67 (green) in CRBOs culture without or with matrigel. Counterstaining of nuclei was done with DAPI (in blue). (O) Quantification of the percentage of KI67+ in full CRBOs or in VZ regions in CRBOs in the absence or presence of Matrigel. (n=5). Data are represented as mean ± SEM. T-test ns= non-significant; *p<0.05; **p<0.01. Scale bars represent 100 μm.

**Figure S2:**
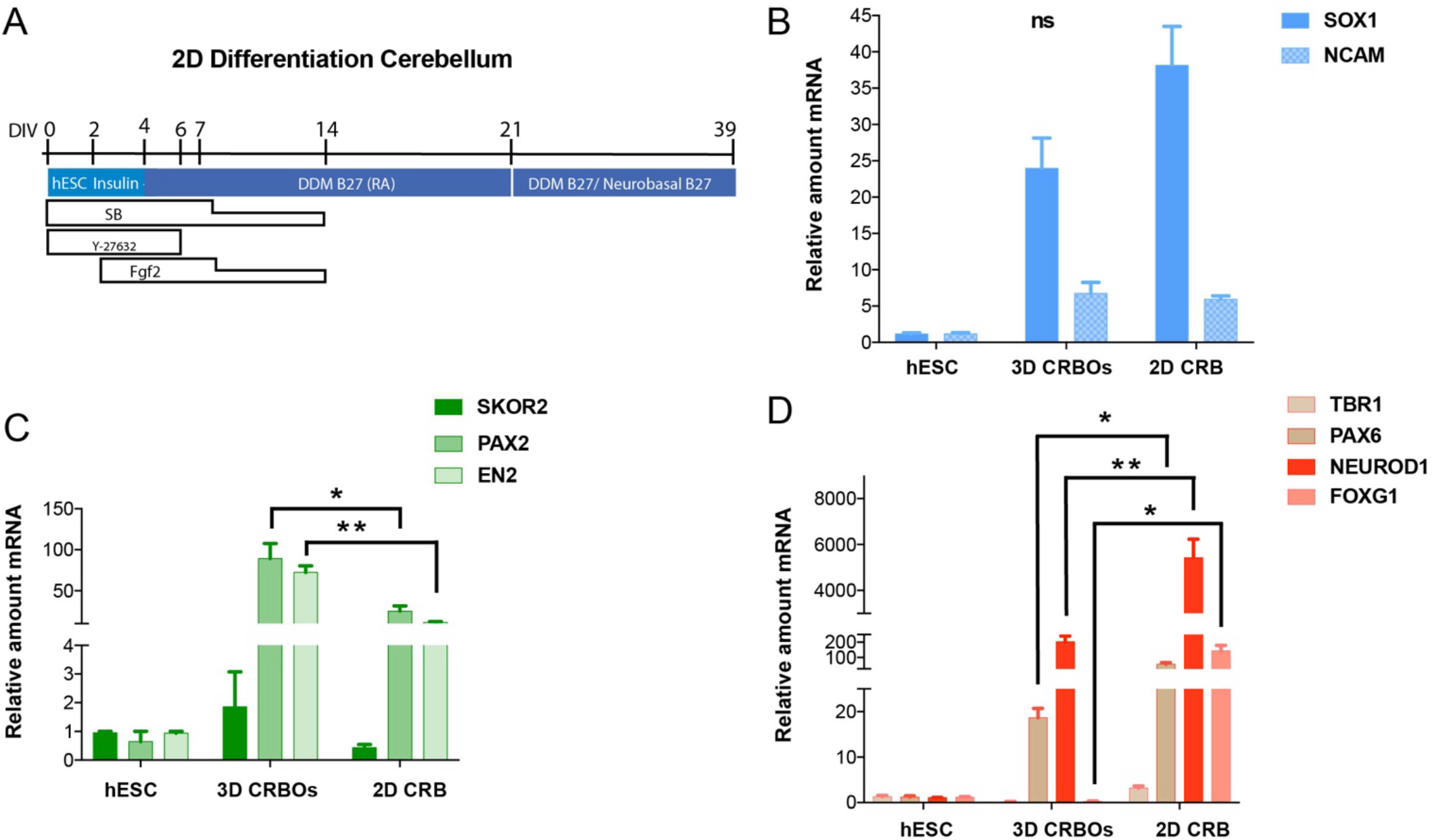
3D CRBOs show restricted caudal cell identity compared to 2D differentiation paradigms. (A) Skin of the protocol for the generation of 2D cerebellar cultures. (B-D) qRT-PCR analysis of the expression of SOX1 progenitor and NCAM neuronal general markers (B), SKOR2, PAX2, EN2 caudal cerebellar-specific markers (C), FOXG1 telencephalic marker (D), and glutamatergic neuronal markers TBR1, PAX6 and NEUROD1 (D) in hESC (day 0), 3D CRBOs and 2D CRB at day 14 *in vitro* (n=2-3). Data are shown as relative amount of RNA compared to the value of hESC as value 1 ± SEM (fold change). One-way ANOVA with Tukey post-tests. ns= non-significant; * p<0.05; **p<0.01.

**Figure S3:**
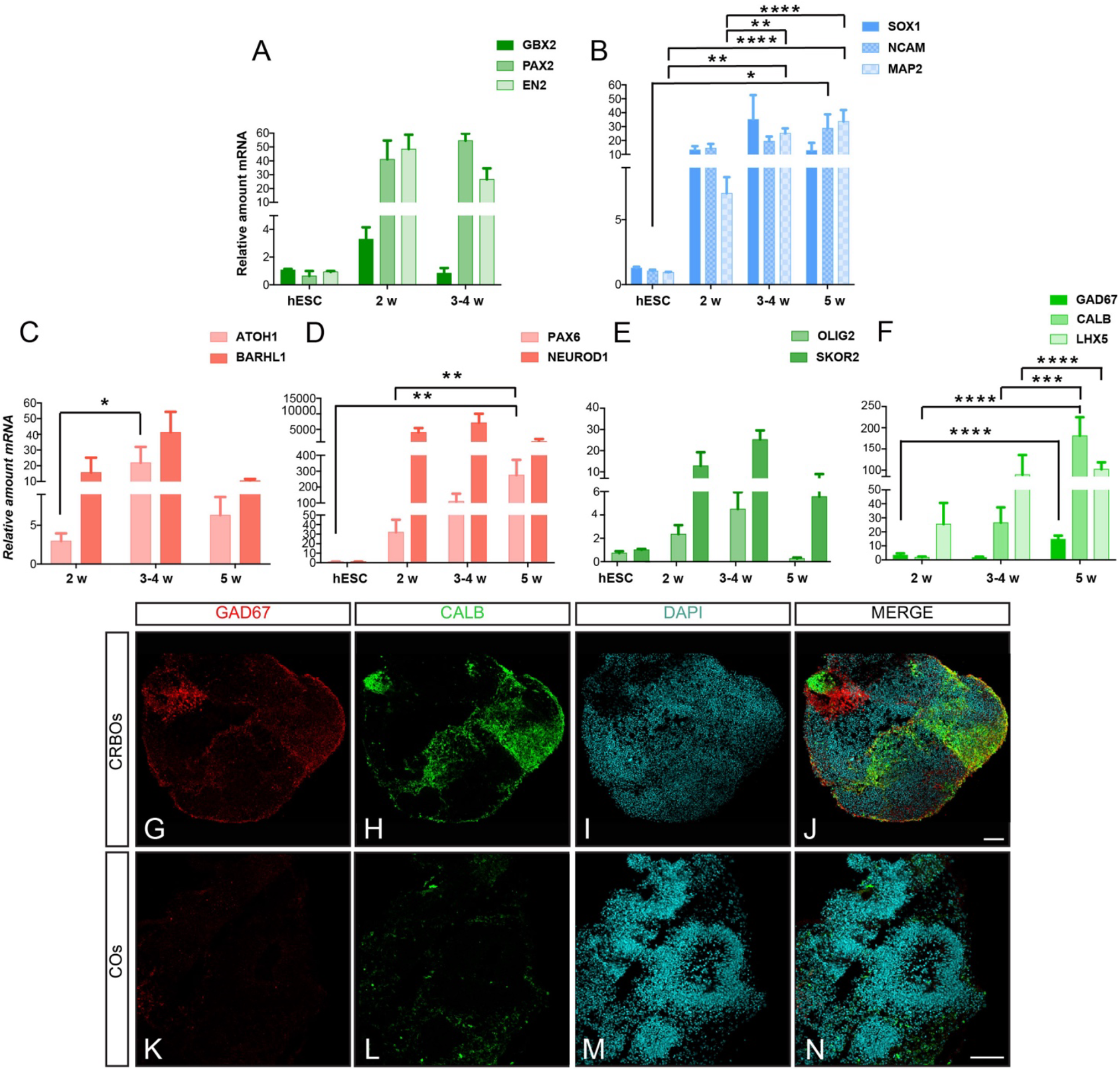
3D CRBOs show restricted caudal cell identity compared to 2D differentiation paradigms. (A) qRT-PCR analysis of the expression of caudal cerebellar regional markers in hESC (day 0) and CRBOs at 2 weeks and 3-4 weeks *in vitro*. (n=2-8). (B) qRT-PCR analysis of the expression of the neuronal progenitor marker SOX1, and the neuronal markers NCAM and MAP2 in hESC (day 0) and CRBOs at 2 weeks; 3-4 weeks and 5 weeks *in vitro*. (n=4-10). (C-F) qRT-PCR analysis of the expression of glutamatergic (C-D) and GABAergic (E-F) neuronal cerebellar markers in hESC (day 0) and CRBOs upon time *in vitro*. (n=3-10). Data are shown as relative amount of RNA compared to the values of hESC or 2 weeks CRBOs as value 1 ± SEM (fold change) (n=5-8). One-way ANOVA with Tukey post-tests. ns= non-significant; * p<0.05; **p<0.01; ***p<0.001; ****p<0.0001. (G-N) Immunofluorescence tile scan images showing the expression of the GABAergic markers GAD67 (red) and CALB (green) in CRBOs (G-J) and COs (K-N) at day 35. Counterstaining was done with DAPI (cyan). Scale bars represent 100 μm.

**Figure S4:**
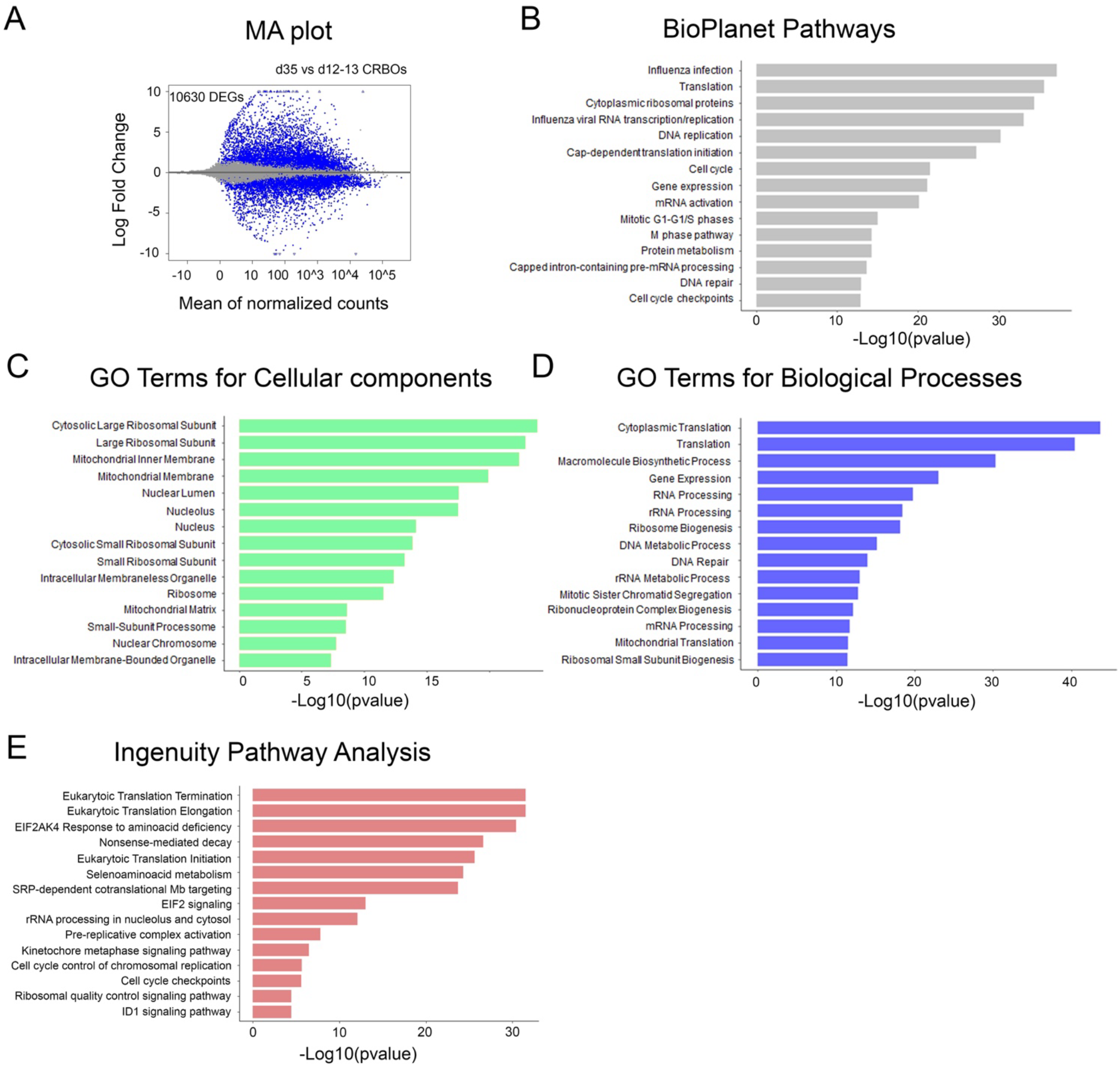
Downregulation of genes associated with translation, splicing, gene expression, cell cycle and mitochondria during the development of hESC-derived CRBOs ki67. (A) MA plot showing DEGs in the CRBOs at 35 days compared to12-13 days in vitro. Each blue dot represents one DEG defined by FDR < 0.05. The x-axis represents the mean of normalized counts and the y-axis the log2-fold change. D12-13=12 and 13 days in vitro CRBOs. d35=35 days in vitro CRBOs. The top 15 results from (B) BioPlanet pathways, (C) GO terms for cellular components, and (D) GO terms for biological processes based on the significance of enrichment are shown. (E) Pathway enrichment analysis over all DEGs (after applying a cut off of absolute fold change > 1) in day 35 compared with day 12-13 in vitro CRBOs using IPA and selecting the downregulated ones based on negative Z score.

## Supplemental Tables

**Supplementary Table 1. List of DEGs from day 35 compared to day 0 *in vitro* CRBOs**. The separate worksheets include total DEGs with FDR < 0.05, upregulated DEGs, downregulated DEGs, and DEGs shared with ataxia-associated genes downloaded from DISGENET database. The table includes ENSEMBL IDs, base mean, log2-fold change, lfcSE, p-value, FDR (padj), and gene symbol for each DEG.

**Supplementary Table 2. List of enriched cell types in CRBOs top20 upregulated DEGs from comparison day 35 versus day 0**. It includes enrichment from DEESCARTES database after filtering for CNS cell types and discarding any other tissue cell types.

**Supplementary Table 3. Ataxia associated genes from DISGENET database**.

**Supplementary Table 4. List of DEGs from day 35 compared to days 12-13 *in vitro* CRBOs**. The separate worksheets include total DEGs with FDR < 0.05, upregulated DEGs, and downregulated DEGs. The table includes ENSEMBL IDs, base mean, log2-fold change, lfcSE, p-value, FDR (padj), and gene symbol for each DEG.

**Supplementary Table 5. Lists of enriched pathways using BioPlanet database in CRBOs DEGs from comparison day 35 versus day 12-13**. Separate worksheets include enrichments for upregulated and downregulated DEGs.

**Supplementary Table 6. Lists of enriched pathways using GO Terms for Cellular Components in CRBOs DEGs from comparison day 35 versus day 12-13**. Separate worksheets include enrichments for upregulated and downregulated DEGs.

**Supplementary Table 7. Lists of enriched pathways using GO Terms for Biological Processes in CRBOs DEGs from comparison day 35 versus day 12-13**. Separate worksheets include enrichments for upregulated and downregulated DEGs.

**Supplementary Table 8. Lists of enriched pathways using Ingenuity Pathway Analysis in CRBOs DEGs from comparison day 35 versus day 12-13**. Negative Z score corresponds to a predicted downregulation of pathways, while positive Z score corresponds to a predicted upregulation of pathways.

**Supplementary Table 9. Lists of predicted upstream regulators of the gene expression changes using Ingenuity Pathway Analysis in CRBOs DEGs from comparison day 35 versus day 12-13**.

## Supplemental Experimental Procedures

### Human Pluripotent stem cell culture

Human ESC H9 line (WiCell, WA09, metadata: female, no disease associated) was cultured using standard procedures for feeder-free cultures (WiCell) and maintained in mTeSR plus medium (StemCell Technologies ref: 100-0276) on geltrex (Thermo Fisher ref: A1413302)-coated plates. H9 cells were passaged using EDTA (Sigma ref:03690) and kept for a maximum of 10-15 passages. The H9 line was authenticated by WiCell upon acquisition and assayed for the expression of pluripotency markers (qPCR and immunofluorescence), and mycoplasm screenings were performed every 3 months. H9 cultures were kept in culture without antibiotics under good laboratory practices. Genomic characterization was performed by copy-number-assay (qPCR CNV) on genomic DNA (Thermo Fisher) from hESC-H9 cells amplified from cryopreserved stocks (passages ∼ 30-40).

### Differentiation of CRBOs from hPSC

Cerebellum brain organoids were developed following a modified version of an already existing protocol (Muguruma et al., 2015). hESC-H9 cell line (Wicell) was dissociated using accutase (Stem Cell Technologies Ref: 7922) and seeded in 96 well U bottom plates (Nunclon sphera Ref: 15396123) by plating 9000 cells per well. Plates were centrifuged 1min at 500 rpm and cells were cultured in DMEM-F12 (ThermoFisher Ref: 31331-093) with 20% KO serum (Fisher Ref: 10828028), supplemented with 1/100 non-essential amino acid (NEAA; ThermoFisher Ref: 11140-035), 1/100 penicillin/streptavidin (Pen/strep; ThermoFisher Ref: 15140122), 1/100 Glutamax (ThermoFisher Ref: 35050-038) and 100µm β-mercaptoethanol (Sigma Ref: 444203) for 14 days. Morphogens were added to the medium for 14 days. From day 0: 10µM SB (SB431542; Sigma Ref: S4317) and 7µg/ml insulin (Sigma Ref: I9278-5ML), chemically defined lipid concentrate (1% Invitrogen). After day 2, 50µg/ml FGF2 (Peprotech Ref: 100-18b) was added. After day 7, the concentration of SB and FGF2 were reduced respectively to 5µM and 25µg/ml. Rock inhibitor (Y-27632 2HC; Bio-Connect Ref: S1049) was provided on day 0 when seeding the cells at 20µM and then at 10µM from day 2 to day 6. From day 14 to day 28, medium was composed of DMEM-F12, 1/100 sodium pyruvate (Na Pyr; ThermoFisher Ref: 11360-039), 1/100 NEAA, 1/100 N2 (ThermoFisher Ref: 17502048), 1/100 Pen/strep, 1/100 Glutamax, 1/150 BSA (BSA fraction V; ThermoFisher ref: 9048-46-8), 1/50 B27 without vitamin A (ThermoFisher Ref: 11500446) and 100µm β-mercaptoethanol. Insulin (7µg/ml) (Sigma Ref: I9278-5ML) and 1% co-lipids were also added to the medium untill day 21. On day 23-25, organoids were transferred in a 6cm dish and placed on orbital shaker (75 rpm). From day 28 to day 35, medium was composed of half DMEM-F12 and half Neurobasal (ThermoFisher Ref: 21103-049) and supplemented with 1/200 Na Pyr, 1/200 NEAA, 1/200 N2, 1/200 Glutamax (ThermoFisher Ref: 35050-038), 1/300 BSA, 1/100 Pen/strep, 1/50 B27 without vitamin A and 50µm β-mercaptoethanol. Medium was changed every 2 days.

### Differentiation of COs from hPSC

Cortical brain organoids are developed following a modified version of an already existing protocol (Kadoshima et al., 2013). hESC-H9 cell line (Wicell) was dissociated using accutase (Stem Cell Technologies Ref: 7922) and seeded in 96 well U bottom plates (Nunclon sphera Ref: 15396123): 9000 cells per well. Plates were centrifuged 2min at 550 rpm and cells were cultured in DMEM-F12 (ThermoFisher Ref: 31331-093) with 20% KO serum (Fisher Ref: 10828028), supplemented with 1/100 non-essential amino acid (NEAA; ThermoFisher Ref: 11140-035), 1/100 penicillin/streptavidin (Pen/strep; ThermoFisher Ref: 15140122) and 100µm β-mercaptoethanol (Sigma Ref: 444203) for 14 days. Morphogens are added to the medium for 14 days: 5µM SB (SB431542; Sigma Ref: S4317) and 3µM IWR1 (Sigma Ref: 681669). Rock inhibitor (Y-27632 2HC; Bio-Connect Ref: S1049) is added on day 0 when seeding the cells at 20µM and then at 10µM from day 2 to day 6. From day 14 to day 28, medium is composed of DMEM-F12, 1/100 sodium pyruvate (Na Pyr; ThermoFisher Ref: 11360-039), 1/100 NEAA, 1/100 N2 (ThermoFisher Ref: 17502048), 1/100 Pen/strep, 1/150 BSA (BSA fraction V; ThermoFisher Ref: 9048-46-8), 1/50 B27 without vitamin A (ThermoFisher Ref: 11500446) and 100µm β-mercaptoethanol. Organoids were transferred to 6cm dishes (Greiner Ref: CLS430196) at day 23-25 and placed on orbital shaker (75 rpm). From day 28 to day 35, medium is composed of half DMEM-F12 and half Neurobasal (ThermoFisher Ref: 21103-049) and supplemented with 1/200 Na Pyr, 1/200 NEAA, 1/200 N2, 1/200 Glutamax (ThermoFisher Ref: 35050-038), 1/300 BSA, 1/100 Pen/strep, 1/50 B27 without vitamin A and 50µm β-mercaptoethanol. Medium was changed every 2 days.

### RNA extraction and sequencing

For all hESC-derived CRBOs samples, RNeasy® Plus Universal Mini Kit (Qiagen, Germany) was used to extract the total RNA according to manufacturer’s protocol. RNA quality was determined by using the Agilent 5300 Fragment Analyzer system. High quality RNA samples with a RQN value above 7 were used for sequencing: 3 samples day 0, 2 samples day 12, 2 samples day 13, 2 samples day 21, 2 samples day 27 and 3 samples day 35. mRNA libraries were generated by the GIGA Genomics Platform (www.gigagenomics.uliege.be) using the TruSeq Stranded mRNA Library Prep kit (Illumina) and sequenced using the Illumina Novaseq 6000 machine to generate paired-end 2×150bp reads.

### RNA-sequencing analysis

Briefly, raw reads were demultiplexed and adapter-trimmed using Illumina bcl2fastq conversion software (v2.20). FastQC (v0.11.9) (Andrews, 2010) was used to assess the quality control of the raw reads. TrimGalore was used to trim a single base at 5’ and 3’ ends of each read and to perform a quality-based trimming after removing poly-G tails. This was followed by mapping using STAR (Dobin et al., 2013). Human reference genome used was GRCh38 release 103. Picard tool was used to validate the BAM files (Adams et al., 2000)and quality was assessed by various QC modules along the pipeline, all were grouped into the final multiQC report. After getting the BAM files, Salmon produced the counts of the reads (Liao et al., 2014). Raw counts were processed using dplyr package in order to generate the reads per kilo base of transcript per million mapped reads (rpkm). Raw count table obtained with Salmon was processed using DESeq2 R package (v1.36.0) (Love et al., 2014) to perform the differential expression analysis and obtain the differentially expressed genes (DEGs). RUVSeq (v1.20.0) was used to remove unwanted variation (batch correction). Enrichment analysis for DEGs was conducted with Enrichr and Ingenuity Pathway Analysis (IPA) (Chen et al., 2013; Krämer et al., 2014; Kuleshov et al., 2016; Xie et al., 2021).

### Cryopreservation and immunohistochemistry of CRBOs and COs

Briefly, hESC-derived CRBOs and COs were immersed in 4% paraformaldehyde for 30 min at 4°C. After two washing steps in PBS 1X to remove excess fixative, hESC-derived CRBOs and COs were immersed in 30% sucrose at 4°C for a maximum of 5 days to prevent future freeze damage. After that, organoids are frozen in OCT blocks using Neg-50 blue (epredia, Ref: 6502B) at -80°C. 14 µm slices were cut in a Cryostar NX70 cryostat, placed on slides (Superfrost Plus Adhesion, Epredia), air dried overnight and stored at -20°C or used directly. After thawing them and wash with PBS (1x), antigen retrieval using DAKO Target Retrieval Solution (1x) (Ref: S1699) was performed. This was followed by three washes with PBS (1x) and permeabilization/blocking for 30 min at RT with PBS containing 3% Bovine Serum Albumin (BSA), 5% Donkey serum and 0.3% Triton-X100. After that, sections were incubated with PBS containing 3% Bovine Serum Albumin (BSA), 1% Donkey serum and 0.1% Triton-X100 with the following antibodies: rabbit anti-OTX2 (Milipore, Ref:AB9566, 1/1000 dilution), goat anti-SOX2 (Santa Cruz, Ref: sc17320, 1/500 dilution), rabbit anti-PAX6 (Ref: PRB-278P Covance, 1/500 dilution), rabbit anti-TBR1 (Abcam, Ref: ab31940, 1/250 dilution), mouse anti-LHX2 (Ref: MA-15834, 1/500 dilution), chicken anti-MAP2 (Abcam, Ref: ab5392, 1/1000 dilution), rabbit anti-OLIG2 (Sigma Aldrich, Ref: ab9610, 1/500 dilution), rabbit anti-CALB (Ref: CB-#38a Swant, 1/2000 dilution), mouse anti-GAD67 (Milipore, Ref:MAB5406, 1/1000 dilution), mouse anti-Ncadherin (Sigma, Ref:C2542, 1/1000 dilution), rabbit anti-Ki67 (Abcam, Ref:ab15580, 1/500 dilution), overnight at 4°C. The next day, primary antibodies are removed and sections are washed three times with PBS (1x) before adding the secondary antibodies: Alexa488 conjugated anti-rabbit (Invitrogen Ref:A21206, 1/500 dilution) Alexa555-conjugated anti-mouse (ThermoFisher, Ref: 831517, 1/500 dilution), Alexa555-conjugated anti-goat (ThermoFisher, Ref: 821432, 1/500 dilution) and DAPI (Sigma Aldrich, Ref: D9542, 1/1000 dilution) for 2h at RT. Slides were then washed twice with PBST and also twice with PBS (1X) for 5 min at RT. After removing PBS, sections were immediately covered by 1-2 drops of DAKO Mounting Gel (Ref: CS703) and a glass coverslip. Slices were dried overnight in the dark and then stored at 4°C. Sections were analyzed under a Zeiss LSM 880 confocal laser scanning with Airyscan microscope at a 20-fold magnification.

### Quantification of the percentage of SOX2+, PAX6+, PAX6+/SOX2, KI67+, LHX2+ cells

Image quantifications were done semi-automatically using ImageJ/Fiji. The threshold used was the following: intermodes for PAX6, default for SOX2, DAPI, and Li for KI67, and default dark for LHX2, with a particles size of minimum 10 μm2. The choice of the threshold value was made in each case by comparing the generated binary picture (white and black) with the original picture (16 bits) and setting the value that would match best the signal observed.

### Quantification of VZ area

The area of the ventricular zone (VZ) was quantified using ImageJ/FIJI software. The VZ was identified based on Sox2 immunostaining. For each one, the VZ region was manually delineated by tracing its boundaries using the segmented line selection tool, ensuring that the defined area encompassed the central portion of the VZ. Measurements were performed on all the area of the organoid.

### Quantification of the CALB+ organoid area

Quantification of CALB+ organoid area was done automatically using a ImageJ/Fiji macro. The first step of the macro is to define the whole area of the organoid. This step is done based on the DAPI signal using the Huang threshold to capture the maximal signal, to have the full organoid recognized as one object. The second step is to define a mask from this object and to apply it on the channel corresponding to the CALB. The threshold used was moments. The third step is to measure the percentage of organoid stained by CALB with the function “%Area fraction” which give this percentage of the organoid stained.

### RT-qPCR

For cDNA synthesis, depending on the sample availability, 1µg of total RNA was used. Total RNA was mixed with the following reagents from Thermo Scientific RevertAid H Minus First Strand cDNA Synthesis Kit (Ref: K1632) to get a final volume of 20 μl: 1 µl of Random Hexamer Primer, 4 µl of 5X Reaction Buffer, 1 µl of RiboLock RNase Inhibitor, 2 µl of 10mM dNTP Mix, 1 µl of Reverse Transcriptase, 12 µl of sample (+DNase/RNase-free water).The reverse transcription was performed at 42°C for 1 hour and stopped by incubating at 70°C for 5 min. After that, cDNA concentration is is 50 ng/µl that is diluted in DNase/RNase-free water to obtain a concentration of 3,125 ng/µl. qPCR reactions were performed using 4 μl (12,5 ng) of diluted cDNA, 0,5 μl of a mix containing 250 nM custom designed primers, 5 μl of GoTaq qPCR Master Mix 2X, and 0,5 μl of DNase/RNase-free water for a total of 10 μl per well. Specificity of the primers was verified previously by detecting a single melting point in the melt curve. qPCR parameters were the following: 95°C for 15 min and then 40 cycles of 95°C for 15 sec, 55°C for 30 sec and 72°C for 30 sec. Gene expression levels were normalized following the 2-(ΔΔCt) method. Briefly, the following formula (ΔCt (target gene)-ΔCt(housekeeping gene)) test condition -(ΔCt(target gene)-ΔCt (housekeeping gene)) reference condition was used to get ΔΔCt. YWHAZ was used as the housekeeping gene. To perform statistics, we used GraphPad Prism 9 and to determine the significance, a two-way ANOVA test was applied to CRBOs samples.

### Primers

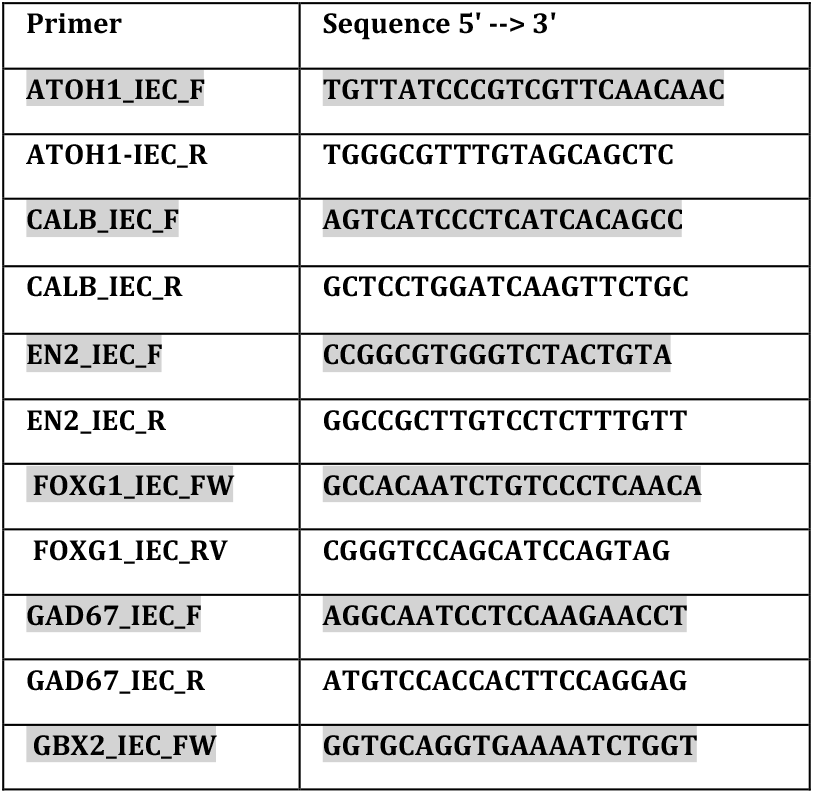
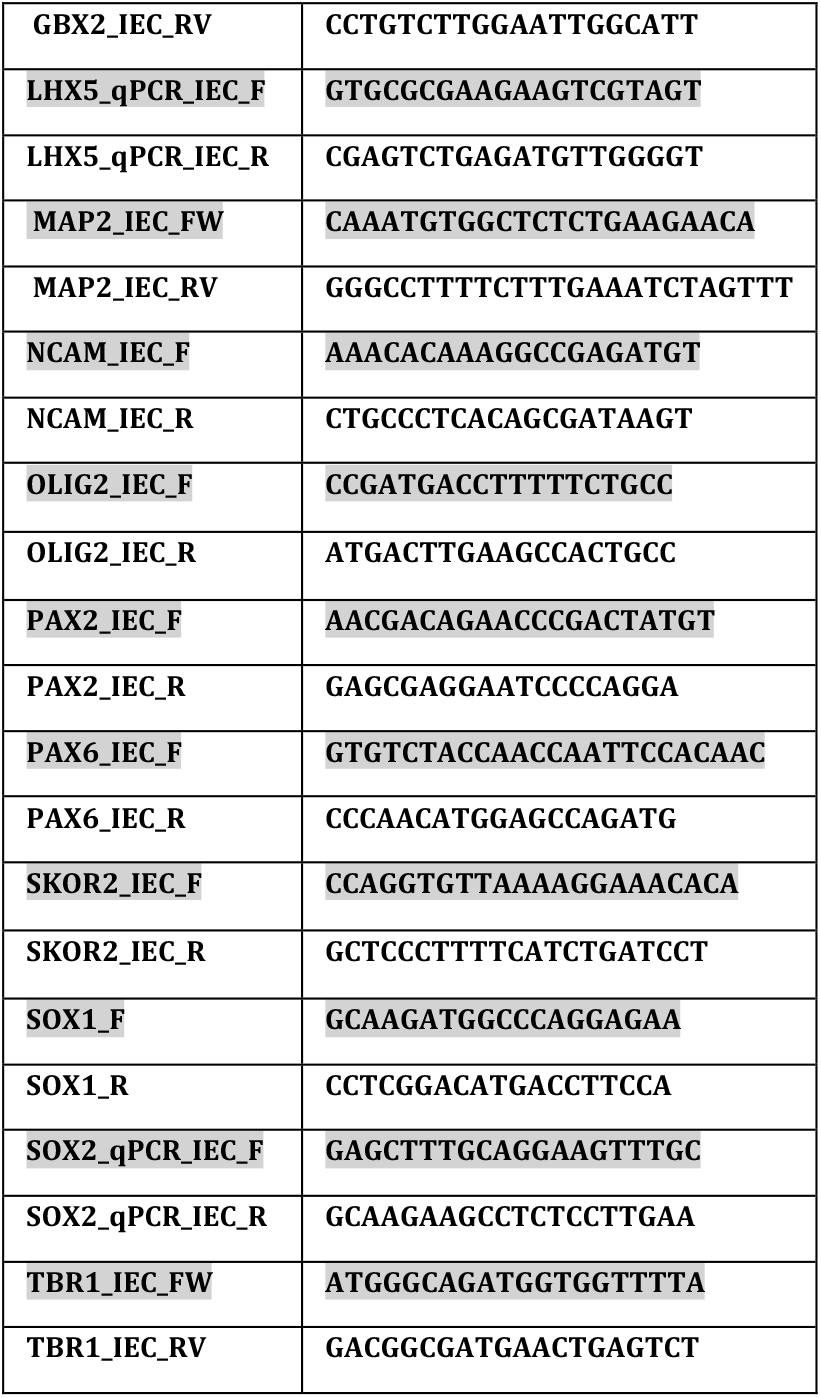

## Notes

### Competing Interest Statement

The authors have declared no competing interest.

